# Constraints on somite formation in developing embryos

**DOI:** 10.1101/645010

**Authors:** Jonas S. Juul, Mogens H. Jensen, Sandeep Krishna

## Abstract

Segment formation in vertebrate embryos is a stunning example of biological self-organisation. Here, we present an idealized model of the presomitic mesoderm (PSM) as a one-dimensional line of oscillators. We use the model to derive constraints that connect the size of somites, and the timing of their formation, to the growth of the PSM and the gradient of the somitogenesis clock period across the PSM. Our analysis recapitulates the observations made recently in ex-vivo cultures of mouse PSM cells, and makes predictions for how perturbations, such as increased Wnt levels, would alter somite widths. Finally, our model makes testable predictions for the shape of the phase profile and somite widths at different stages of PSM growth. In particular, we show that the phase profile is robustly concave when the PSM length is steady and slightly convex in an important special case when it is decreasing exponentially. In both cases, the phase profile scales with the PSM length; in the latter case, it scales dynamically. This has important consequences for the velocity of the waves that traverse the PSM and trigger somite formation, as well as the effect of errors in phase measurement on somite widths.

## I. INTRODUCTION

A particularly striking example of biological self-organisation is that of segmental patterning in vertebrate embryos. During somitogenesis in vertebrate species, somite segments, the precursors of vertebrae, form periodically as the embryo elongates. In mice, chick, and zebrafish embryos, cells in the presomitic mesoderm (PSM) behave like a population of coupled oscillators. Expression of many genes oscillate in each cell, and cells coordinate their oscillations such that kinematic waves of gene expression travel from the posterior end of the PSM to the anterior. The arrival of each wave at the anterior end is correlated with the formation of a new somite [1–4]. In this paper, we investigate the constraints that connect these waves to the somite width and the gradient of oscillation periods across the PSM.

Several genes are known to oscillate in the PSM of vertebrates, most importantly those in the Notch, Wnt and FGF pathways [5]. The period of oscillations often depends on the position of the cell along the antero-posterior axis. There is a region in the tail bud where all cells oscillate synchronously with a time period characteristic of the species, which can range from ≈30 min for zebrafish to ≈2 hrs in mice. The oscillations slow down as one moves from the posterior end of the PSM (right after the tail bud) to the anterior end [1, 4, 6]. In mice, this “period gradient” is linear – see [4], who find that the posterior-most cells oscillate with a period ≈ 130 min, linearly increasing to 25%-30% higher for the anterior-most cells.

As mentioned above, examining how the oscillations develop over time revealed travelling kinematic waves of gene expression that move from posterior to anterior. For instance, Lauschke et al. [4] report that the position of peak levels of LuVeLu, a Notch signalling reporter, moves from posterior to anterior in ex-vivo cultures of mouse PSM cells (so called mPSMs), with a velocity that depends on the length of the mPSM [4]. Similar waves are observed in a reporter for the oscillating gene *her1* in zebrafish [3]. An important difference between these species is that in zebrafish, several waves can simultaneously co-habit the PSM [3], whereas experiments on mPSMs have found maximally one wave existing at a time [4]. However, in both cases, as well as in other species, the formation of the next somite is coincident with the arrival of a wave at the anterior end, in the vicinity of the previous somite. The mechanism that triggers the formation of a new somite is still a matter for debate. It was thought for years to be the classic clock-and-wavefront model [7], but this theory has recently been challenged. Cotterell et al. [8] combine theory and experiments to suggest that, in chick embryos, formation of new somites might be caused by a reaction-diffusion mechanism in the anterior PSM that interacts with the oncoming wave, while Sonnen et al. [9] suggest that interactions between two different oscillating pathways may be what triggers somite formation in mice.

Regardless of the mechanism, some interesting observations have been made about the periodicity of somite formation and scaling of the somite widths. In mPSMs, the formation of a new somite was found to occur when 2*π* of phase (i.e., one full wave) was spanning the PSM. That is, when a wave reached the anterior end, and a new somite formed, the next wave was just setting out from the posterior end. Furthermore, each new somite consisted of the anteriormost cells that contained 21% of the total phase difference across the PSM, irrespective of the length of the PSM at that time [4]. Although many different aspects of the coupled oscillating cells in the PSM have been investigated theoretically, ranging from models of global wave patterns and morphogen gradients, to models of the underlying biological clocks, and the effect of couplings on defect-free patterning [8, 10–20], nothing is known about the measurable consequences of such phenomenological observations about the phase of the cells in the PSM. In the present paper, this is what we seek to illuminate. A second goal of our work is to understand the interplay between such oscillations (and travelling waves) and the growth of the PSM. Across species, the PSM is known to elongate at the posterior end as the tail bud extends. The length of the PSM is determined by a combination of this growth at the posterior end, and shrinkage at the anterior end as new somites are formed. During somitogenesis, the PSM length typically initially increases, then may remain steady for a duration and finally decreases (indicating an eventual decrease in the growth rate at the posterior end). We examine how the period gradient, growth of the PSM, and shrinkage due to somite formation combine to affect the phases of oscillating cells, and what quantitative constraints this places on the somite widths and the timing of their formation.

The rest of the paper is structured as follows. In Section II, we introduce our model and key assumptions. In Section III A, we show that the period gradient, the total phase difference across the PSM at somite formation, the growth rate of the PSM and the width of the new somite cannot be independent of each other. We explicitly derive the mathematical constraint that connects these quantities and, in Section III A 1, show that experimental measurements from mPSMs match this constraint. Section III B calculates the phase profile across the PSM in the specific situation where the PSM length is in steady-state, i.e., it is shortened by somite formation at the same rate as it grows at the posterior end, and Section III C calculates the constraints on somite widths that exist in a PSM with steady-state length. Our analysis provides explicit predictions for how the phase of a cell should depend on the antero-posterior location of that cell in wild-type embryos that abide by these constraints (Section III B), and for the expected change in somite widths in an experiment that would perturb the period gradient (Section III D). Finally, we examine the case where PSM growth is arrested, similar to the end of somitogenesis, and make predictions for how somite widths and the PSM length change with time in this case (Section III E). Section IV discusses the experimental predictions stemming from our analysis and speculates on the implications for somitogenesis.

## II. THEORETICAL FRAMEWORK FOR ANALYZING THE PHASE OF THE OSCILLATING CELLS IN THE PSM

We focus on the phase of the oscillation in cells rather than the full waveform of gene expression levels. That is, we associate to each cell a single dynamical variable taking values between 0 and 2*π* representing the phase of the somitogenesis clock in that cell, and a time period that sets the rate of change of the phase. In doing so, we make the implicit assumption that varying the time period simply scales the oscillation waveform without changing its shape otherwise. This seems to be consistent with experimental data (e.g., see Fig. 2 in [3]) and is what allows us to characterize each cell by a single variable, its phase, and a single parameter, its time period, that controls how quickly the phase changes. Further, we simplify the PSM into a 1-dimensional line of cells since the spatial periodicity in somite formation is along the posterior-anterior axis.

**FIG. 1.**
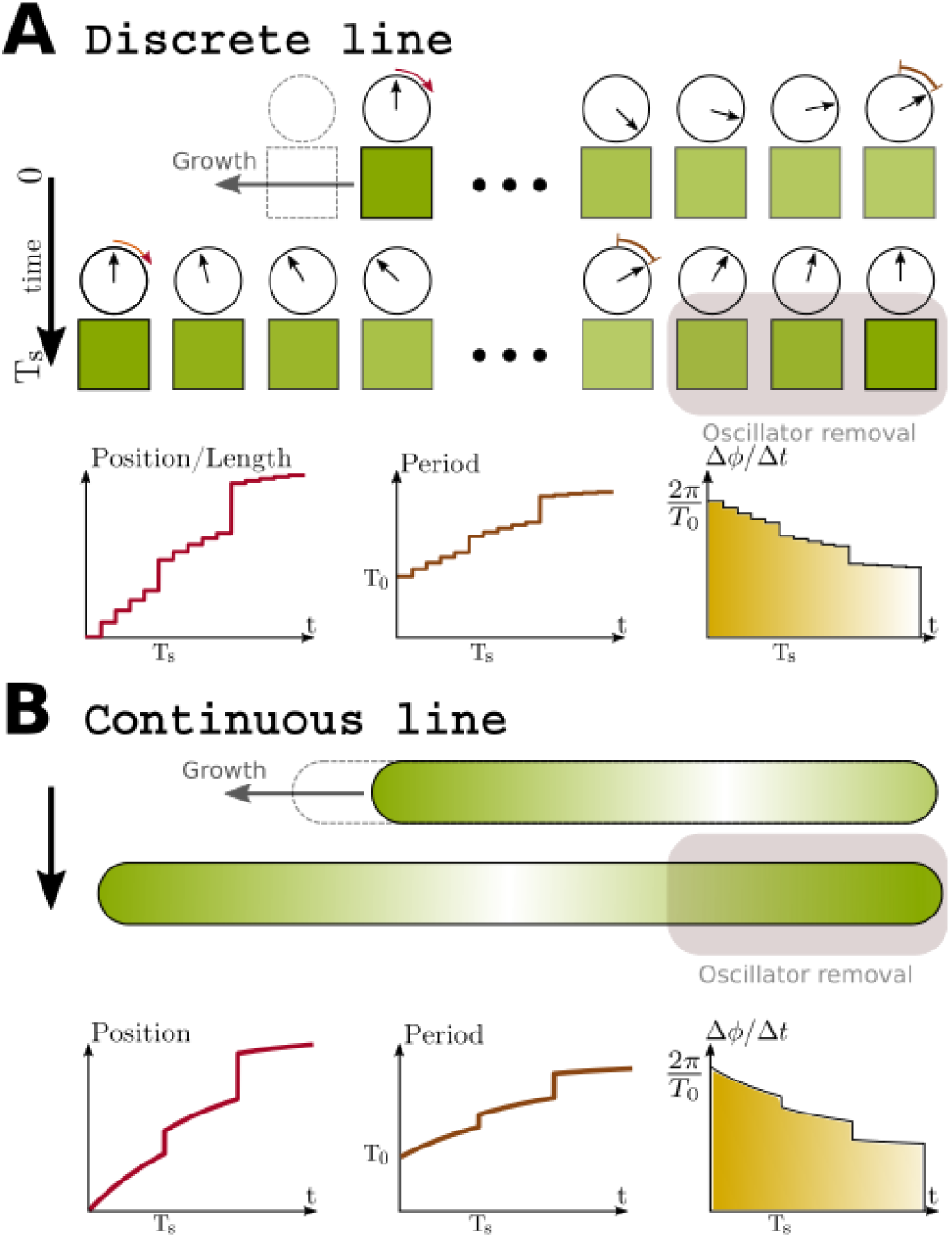
Illustration of the idealised PSM. **A:** Discrete system. The PSM is approximated as a finite number of oscillators on a line. The phase of each oscillator changes according to a position-dependent oscillation period *T* (*x*). The posterior-most cell is located at *x* = 0, while the anteriormost cell is at *x* = 1. The relative position of an oscillator changes as cells are gradually added to the posterior (with period *T*_*g*_), and removed (with period *T*_*s*_) in chunks from the anterior end. As time progresses, each cell effectively moves toward the anterior; the three insets show, as a function of time, the relative position, oscillation period and change of phase per time of a cell which is initially located at the posteriormost position. **B** The same as in A, but in a PSM where the relative position *x* does not take a finite number of discrete values, but is taken to be continuous *x* ∈ [0, 1]. This approximation is justified when the number of cells in the PSM is large.

**FIG. 2.**
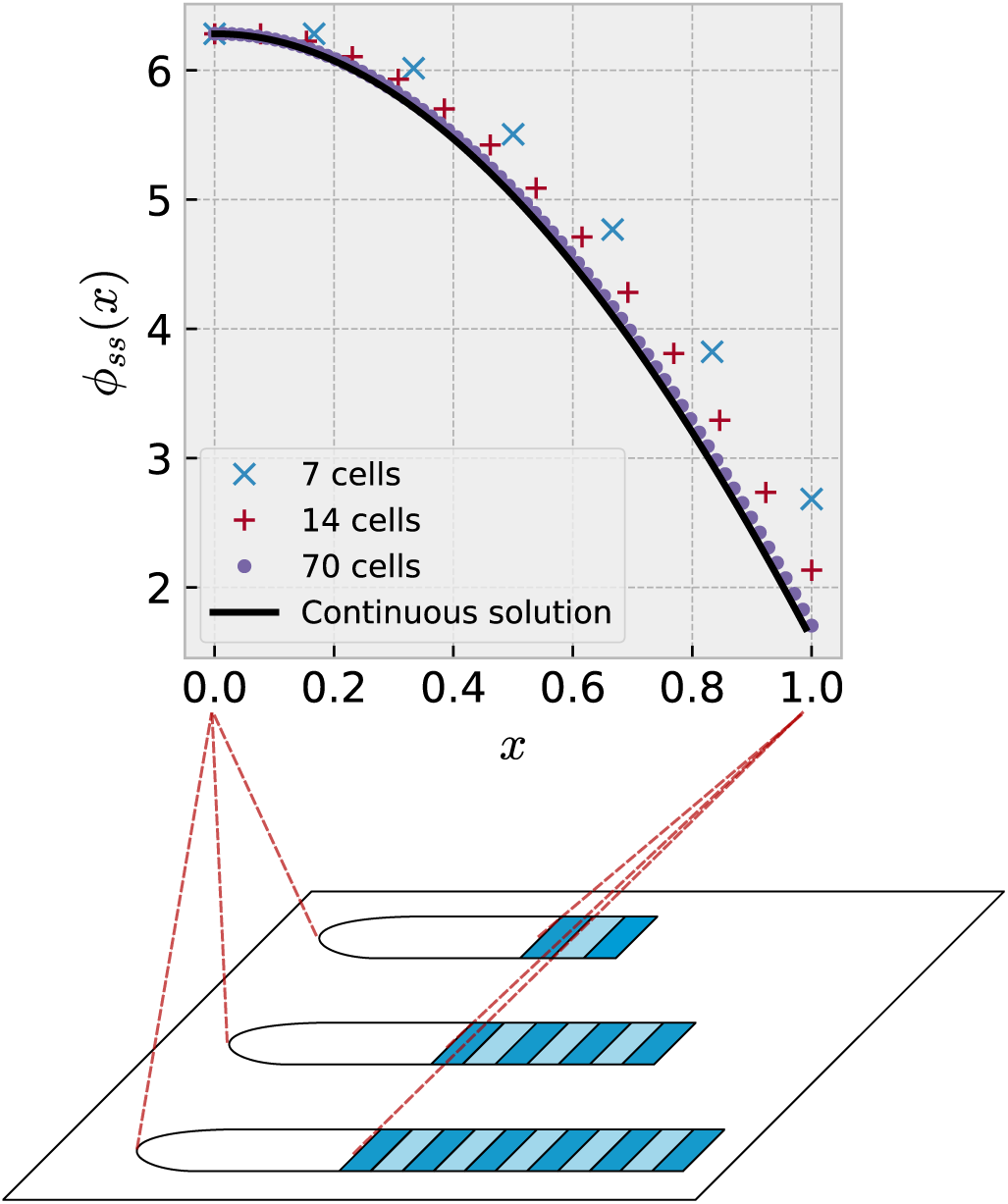
Steady-state phase profile of a PSM of constant length, *ϕ*_*ss*_(*x*). The black curve shows the steady-state phase profile in the continuum limit (calculated from Eq. (16) in Supplementary section 2), when we choose *T*_*s*_ = *T*_0_, *λ* = 0.266, Φ_before_ = 2*π*, and 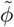 and *T*_*g*_ are chosen such that the length of the PSM varies in a sawtooth manner as follows: ℒ (*t*) = *L*_0_(1 + (*t* mod *T*_0_)/(7*T*_0_)). Note that due to the freedom to choose units of time and length, *ϕ*_*ss*_(*x*) will not depend on what specific values we choose for *T*_0_ and *L*_0_. Also plotted are the steady-state phase profiles calculated for PSMs consisting of a finite number of cells (symbols correspond to PSM lengths after somite formation, *N* = 7 (blue ×), 14 (red +) and 70 cells (purple •.) These profiles are calculated for the case where *T*_*s*_ = *T*_0_, *λ* = 0.266 and *T*_*g*_ and 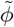 are chosen such that the PSM length varies in a sawtooth manner as *N* (*t*) = *N* + └*t/T*_*g*_ ┘mod *N/*7. We numerically approximate the phase profiles the discrete calculation of Supplementary section 1 would produce for these parameters, by simulating a discrete PSM with length varying as above and updating the phases of each oscillator in time according to Eq. (7) in Supplementary section 1 until steady-state is achieved. Here, Φ_before_ is determined by the remaining parameters, and as seen in the plots, converges to the value obtained in the continuum calculation when *N* becomes large. Note the concave shape of all the phase profiles plotted. In Section B1 and Supplementary Section 6, we show analytically that this concave shape is robust to changes in parameter values and holds for all increasing period gradients, linear or nonlinear.

Thus, the system we consider, see Fig. 1, consists of a 1d line of cells, each associated with a phase and a time period, pictorially represented by a clock face with the clock hand showing the current value of the phase. As observed in embryos, new oscillators are frequently added at the posterior end of the PSM and, when a new somite is formed, oscillators are removed from the anterior end of the PSM. Thus, we allow cells to be added to the posterior end periodically every *T*_*g*_ time units (1*/T*_*g*_ is thus the PSM growth rate)[21], and removed from the anterior end whenever a somite is formed. The evolution of the phase of each cell depends only on the time period of that cell, which in turn depends only on the location of the cell on the line. Thus, the period of a cell may change as addition or removal of cells changes the relative distance of the cell from the posterior end of the line. Travelling waves can occur in this setup. For instance, if the periods of all cells were identical, but the phases initially decreased progressively from 2*π* at the posterior (left) end to 0 at the anterior (right) end, then over time one would observe that the location of phase 2*π* (or 0) would move from left to right, corresponding to a travelling wave moving from posterior to anterior (see Fig 1). In this purely illustrative scenario, the speed of the wave would depend only on the initial phase differences between adjacent cells but, in general, the periods may be different for different cells, in which case the speed of the wave would depend on the period gradient as well as the phase differences.

### A. Key assumptions

We make the following assumptions regarding the phases and periods that characterize the oscillations of each cell:

A. Cells oscillate with a time period *T*_0_(1 + *xλ*), where *x* is the location of the cell relative to the posterior end, normalized to the total length of the PSM (thus *x* ∈ [0, 1]), and *T*_0_ is a species-dependent base time period.
B. A new cell that is added to the posterior end, whenever the PSM grows, is assigned a phase identical to its immediate neighbor, the cell that was until then the posterior-most PSM cell. Subsequently, of course, the phases may start to differ as the two cells will have different time periods.

Assumption (A) posits a linearly increasing period gradient, similar to observations in mPSMs [4], as discussed earlier. In Section III B we show that our key results hold for any increasing period gradient, but for now we assume that the period gradient is linearly increasing. Assumption (A) also implicitly assumes that as new cells are added and removed, due to growth and somite formation, the morphogen gradient determining the periods is quickly reset in such a way that the new posterior and anterior ends retain their periods, *T*_0_ and (1+*λ*)*T*_0_ respectively. This is justified by observations in real embryos and ex-vivo cell cultures in mice: In embryos, the time period of somite formation, which also coincides with the time period of the posterior-most cell, is found to be stable at ≈ 2 h between days 8 and 13.5, during which time more than 60 somites are formed [22]. In ex-vivo experiments, the posterior period has been found to be stable at ≈ 130 min while the tissue was shortening periodically, and other cells slowed down their oscillations as they moved towards the anterior of the colony, ending up with periods of length ≈ 170 min [6] when they were located at the anterior end of the PSM. Note that, when a new somite is formed, this implies that the period gradient (in real length units) becomes steeper. If the phase differences between oscillators in such a resetting were not altered too much, then such a steepening of the gradient should result in slower travelling waves in the smaller PSM. This matches experimental observations [4]. Assumption (B) seems reasonable given that cells in the tailbud and the posterior end of the PSM show stable synchronized oscillations.

Note that we do not explicitly include inter-cellular coupling between the phases of the adjacent cells. However, we do implicitly take into account effects coupling would have on the time periods of cells because we use the empirically observed time period gradient. For a line of coupled oscillators, the time period of each oscillator will be determined both by external factors (e.g., morphogen gradients) that affect the natural (uncoupled) time period, as well as the coupling to adjacent oscillators. A sufficiently strong coupling between adjacent oscillators in a 1-dimensional line can lead to complete synchronization of all the oscillators even if they had substantially different uncoupled time periods, so the coupling must be relatively weak to allow the time period to vary across the PSM. We therefore proceed with the assumption that such a weak coupling would have little effect on the dynamics of the phases of the cells beyond maintaining the period gradient, and perhaps also mitigating the effects of noise on the phases. Hence, for our purpose it is sufficient to include the coupling only implicitly by using the empirically observed period gradient.

With the assumptions mentioned above, we will attempt to obtain and study phase profiles *ϕ*(*x*) that are in steady state. By steady-state, we do not mean that *ϕ*(*x*) is time independent, but rather that *ϕ*(*x*) is the same, modulo 2*π*, at corresponding times between somite formation (for example, right before, or right after, a somite forms). This means that the phase profile exhibits what has been termed ‘dynamical scaling’ in the literature [23], i.e., as the PSM changes in length the pattern of oscillations across it scales correspondingly. We will impose the constraint that new somites are formed from the cells that contain the anteriormost 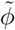 of phase. We shall refer to 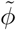 as the phase width of the somite. This constraint, and the scaling of *ϕ*(*x*) with PSM length, are the key observations of recent experiments [4], the consequences of which we set out to explore.

## III. RESULTS

### A. The period gradient constrains the somite width and vice versa

Let *ϕ*(*t, x*) denote the phase of a cell at time *t* and location *x*, where *x* ∈ [0, 1] is the distance from the posterior end, normalized by the PSM length. Let Δ*ϕ*(*t*) ≡ *ϕ*(*t, x* = 1) - *ϕ*(*t, x* = 0) denote the total phase difference across the PSM at time *t*. Assumptions (A) and (B) imply that between somite formations Δ*ϕ*(*t*)) increases linearly in time:

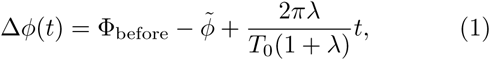

where we assume the previous somite formed at time *t* = 0 and left a total phase difference of 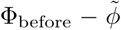 across the PSM just after somite formation 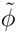 is the phase-width of the somite, described previously, and Φ_before_ is the total phase difference across the PSM before the somite is formed). If the phase profile is in steady-state just before every somite formation event, then it must be that the total increase in Δ*ϕ* between somite formations must exactly match 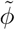, i.e.:

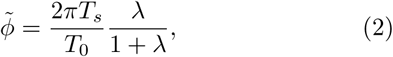

where *T*_*s*_ is the time at which the next somite forms. Because we are considering a steady-state, the phase of the anterior-most cell of the PSM must also be the same (modulo 2*π*) before each somite formation. Therefore *T*_*s*_ must be a multiple of *T*_0_, and we obtain:

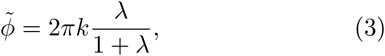

where *k* is a positive integer. Thus, assuming steady state implies that the slope of the period gradient, *λ*, and the phase-width of the somite, 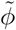, cannot be independent. Note that here we only assume that Δ*ϕ* is in “steady-state” – this does not necessarily imply that the PSM length is a constant before each somite formation. Assuming that the length is a constant imposes additional constraints. Note also that the PSM growth rate does not appear in Eq. (3). Its role emerges in determining the width (as opposed to the phase width) of somites. Both these issues will be explored in Section III B.

#### 1. Comparison with data

In the mouse PSM, the period gradient has been measured in ref. [6] along with the phase width of the newly formed somites [4]. They find that *λ* ≈ 0.275, 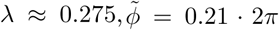 and *T*_*s*_ = *T*_0_ = 130 min. All numbers are not provided with experimental error bars in ref. [4], but even with as low as 5% error, using *λ* = 0.275 in Eq. (3) gives 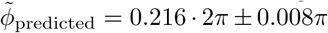, while using 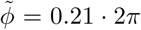 in Eq. (3) gives *λ*_predicted_ = 0.266 ± 0.017. Either way, the experimental observations are consistent with Eq. (3).

### B. When PSM length is constant steady-state phase profile is concave in shape

As mentioned, Eq. (3) does not assume that the length of the PSM right before (or after) each somite formation is a constant. Adding the assumption that the PSM length is also in steady-state allows us to calculate not just Δ*ϕ* but also the entire steady-state phase profile, which we will denote *ϕ*_*ss*_(*x*). Supplementary sections 1 and 2 show this calculation both for the continuum limit, where the number of cells in the PSM is assumed to be infinite, and for the discrete case where the number of cells are finite.

Fig. 2 shows the steady-state phase profile obtained from our calculations when *T*_*s*_ = *T*_0_, *λ* = 0.266, 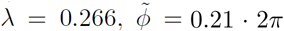, PSM lengths just after somite formation are *N* = 7 (blue ×), 14 (red +) and 70 cells (purple •), and *T*_*g*_ is chosen such that we obtain maximum PSM lengths of *N* (1 + 1/7). These parameters, based on the observations of [4]result in a phase profile with a *concave* shape. The curve is concave both immediately before and after somite formation, since somite formation amounts to removing the anteriormost part of the pre somite-formation curve, and ‘stretching’ the remaining part to cover the full interval [0, 1], neither of which changes the concavity.

When *N* is large enough, the phase profile is indistinguishable for different *N*, which means that the PSM exhibits scaling – the entire somitogenesis pattern scales with the real length of the embryo but does not change in structure otherwise. The calculation for large *N* also matches our continuum calculation for a PSM with infinitely many oscillators, which is shown by the black, continuous line in Fig. 2 (see Fig. 1B for a schematic for the continuous approximation of the PSM).

A testable prediction from our model is that the steady-state phase profile is not linear, but concave in shape. This has consequences for the speed of the travelling waves and reduces the influence of errors in differentiation decisions on somite size, which we will return to in the Discussion. The concave shape is in fact a robust feature of the steady-state phase profile whenever the PSM length is in steady-state, the growth rate is constant and the time period of cells *T* (*x*) is an increasing (linear or non-linear) function of *x*. We demonstrate this in the next section.

#### 1. Concavity is a robust property of the steady-state phase profile for any increasing period gradient

That the steady-state phase profile must be concave in shape for any increasing *T* (*x*), can be seen from the following general argument.

Suppose that the PSM consists of a very large number of cells, so we can use the continuous variable *x* ∈ [0, 1] to describe a cell’s position relative to the posterior (at *x* = 0) and the anterior end (at *x* = 1). Let *T* (*x*) be the period gradient of the PSM, and let this be increasing from posterior to anterior. Suppose that one cell has initial position *x*_0,first_ = *x*\*, and another has initial position *x*_0,second_ = *x*\* + *ϵ*, where 0 ≤ *x*\* < 1, and 0 < *ϵ* ≪ 1. Let us assume *t* = 0 to be immediately after somite formation, and let the phase difference between the two cells at this time be *δϕ*_*ϵ*_ = *ϕ*(*x*) *ϕ*(*x* + *ϵ*) > 0. We now examine how the phase difference between these cells changes between *t* = 0, and the time following the next somite formation at *t* = *T*_*s*_. The change in phase difference between the two cells in this time period is

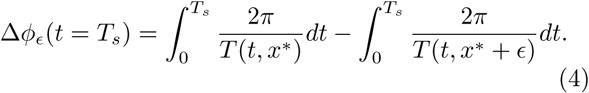

Now, since *ϵ* ≪ 1, we expand the fraction[24]

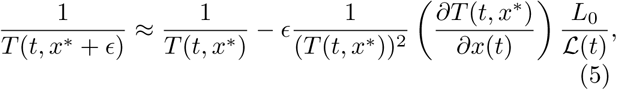

where *L*_0_ is the length of the PSM at *t* = 0, and ℒ (*t*) is the length of the PSM at time *t* ≥ 0. Inserting this expression in Eq. (4) yields,

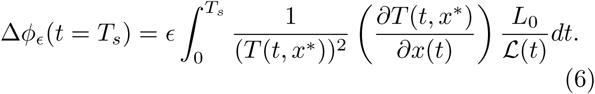

Since *T* (*x*) is increasing and positive, and since ℒ (*t*) is positive and increasing between successive somite formations, Δ*ϕ*_*ϵ*_ > 0. This means that the phase difference between the two cells increases between the two successive somite formations. The phase difference is the same after the somite formation at *t* = *T*_*s*_, and because the PSM length is in steady state, the difference in position between the two cells is still *ϵ* after the somite formation at *t* = *T*_*s*_. The convexity or concavity of the phase profile is determined by the its second derivative – a decreasing, concave function has a negative second derivative, while the second derivative is positive for a decreasing, convex function. An alternative formulation of this is that a decreasing, concave function decreases faster at larger values of the variable it is plotted against, while a decreasing, convex function decreases slower for larger values of the variable. We shall use this formulation to show generality of the phase profile concavity.

The phase profile gradient between the cells at their initial position is *δϕ*_*ϵ*_*/ϵ*, and the phase profile gradient between the cells at their final position is (*δϕ* + Δ*ϕ*_*ϵ*_)*/ϵ*. Calculating the ratio yields

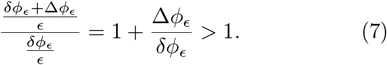

From this we conclude that the steady-state phase profile decreases faster as *x* is increased; or equivalently, the steady-state phase profile is concave.

### C. Constraint on phase differences when PSM length is constant

Now that we have calculated *ϕ*_*ss*_(*x*) we can ask what is the phase difference across the PSM in this state. Following exactly the same argument as in Section III A, it must be true that 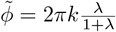 However, in this case we can also derive the actual width of the somite, i.e., the number of cells removed from the anterior end, which must equal the number of cells added between somite formations, *T*_0_*/T*_*g*_. Since the steady-state phase profile scales with respect to the PSM length right after somite formation, *N*, it is of interest to calculate the fractional width of the somites *β* ≡*T*_0_/(*NT*_*g*_) (i.e., *β* is defined as the width of the somite divided by the length of the PSM just *after* somite formation). Just before somite formation, this fractional width must satisfy:

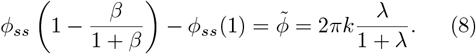

Similar to Eq. (3), this is a constraint between the fractional somite width *β*, the period gradient and the parameters that determine *ϕ*_*ss*_(*x*), namely, *T*_0_, Φ_before_, and 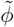. See Supplementary Section 5 for more details on how the phase width, 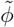, can be converted to the fractional width of the somite, *β*, using this constraint.

Fig. 3 shows a heatmap of this constraint, derived from our continuum calculation, when *T*_*s*_ = *T*_0_ and 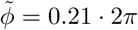. The colours show the value of Φ_before_ that satisfy the constraint Eq. (8) for different values of *β* and *λ*. This heatmap is another prediction of our analysis. Qualitative features that should be experimentally observable include the following: The phase difference between posterior and anterior right before somite formation, Φ_before_, (i) decreases with somite size *β* (for fixed *λ*), (ii) increases with *λ* (for fixed *β*), and (iii) the line *β* ≈*λ/*2 corresponds to the special case Φ_before_ = 2*π*. Prediction (iii) suggests that any change in the period gradient in the mPSM ex-vivo cultures should result in exactly the same change in the fractional width of the somites. Moreover, our calculations predict that this linear relationship depends on there being exactly one wave spanning the PSM at a time. If the system exhibited multiple waves, say Φ_before_ = 4*π* corresponding to two waves, then the relationship between *β* and *λ* would be non-linear and not linear.

**FIG. 3.**
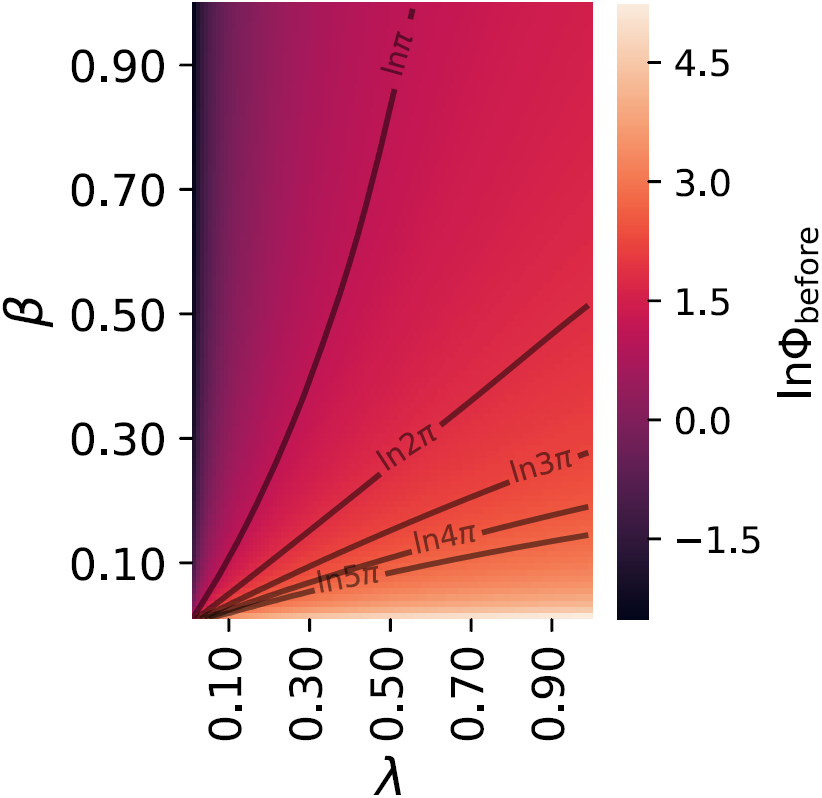
Heatmap of logarithm of the total phase difference across the PSM just before somite formation, Φ_before_, in steady-state phase profiles when the PSM length is constant, as given by the analytical continuum calculation of the constraint Eq. (8). The phase difference is 2*π* on a line *β* ≈ *λ/*2.

### D. Variation of somite width caused by perturbing the period gradient

Assuming that the general constraint of Eq. (3) holds in embryos that are perturbed in various ways, our framework makes specific predictions for the effect of such perturbations. A perturbation that could be feasible to implement experimentally, for instance by affecting the Wnt or FGF gradient in the PSM, would be to change the period of *all* cells by the same additive amount *ξT*_0_. Equation (3) would then become:

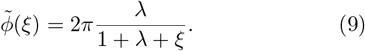

In Fig. 4*A* we show how the somite phase width varies with *ξ*, assuming all other parameters remain the same. Using our analytical calculation in the continuum limit of a PSM of constant length, we can convert the predicted phase width of somites to an actual fractional width (as described above and in Supplementary Section 5). The result is shown in Fig. 4*B*. Thus, we predict that increasing (decreasing) the period of the cells in this manner would decrease (increase) both the phase width and actual width of the somites. Generally, the fractional width of somites, *β*, will be a non-increasing function of *ξ* whenever the steady-state phase of cells decreases from posterior to anterior.

**FIG. 4.**
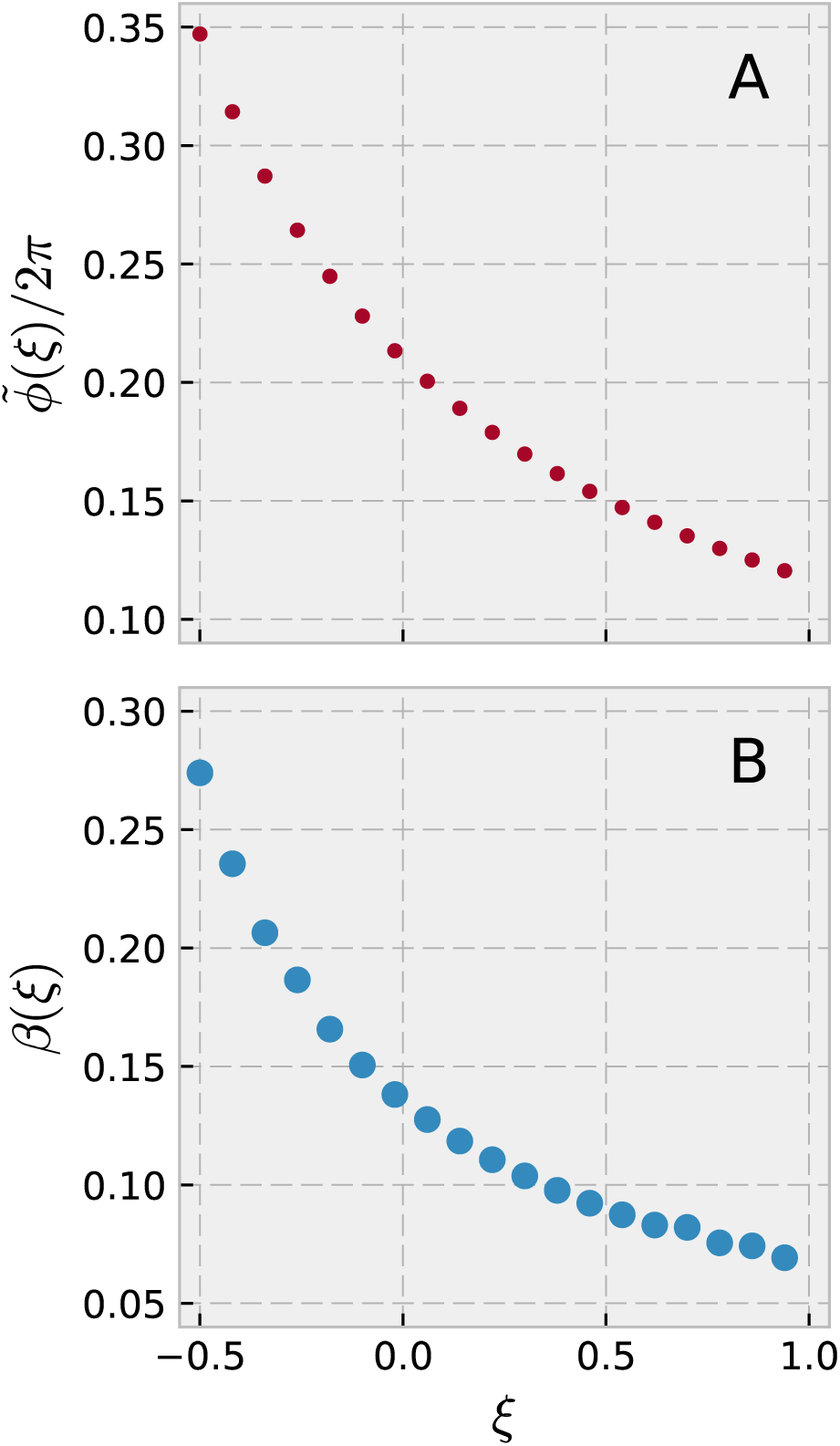
Effect of perturbations on somite widths. Assuming *T*_*s*_ = *T*_0_, and that the constraint expressed in Eq. (3) holds under perturbation of periods in the PSM, we predict that perturbing all periods by an additive amount *ξT*_0_ will alter somite width. **A** The phase width of somites (red dots) will decrease with *ξ*, and is described by Eq. (9). **B** In a PSM of constant length, phase width can be mapped to the actual spatial width of the somite using the continuum solution plotted in Fig. 2. We find that this spatial width also decreases with *ξ* as shown by the blue dots.

### E. Physical somite size and convexity of phase profile in PSMs with no growth

Finally, we consider a case where after the system has reached the steady-state described above, new cells stop being added to the posterior part of the PSM but cells continue to be cut off from the anterior end when new somites are formed. This approximates the very end of somitogenesis (although there the rate of addition of new cells decreases continuously over time rather than falling abruptly to zero). When no new cells are added to the PSM, but the phase across the PSM is in steady-state, we find (see Supplementary section 3) that the length of the PSM of course decreases with time, shrinking by a constant multiplicative factor after each somite formation, which results in an exponential decrease of PSM length with time[25]. Nevertheless, our calculations (see Supplementary section 3) show that the phase profile can attain a steady-state. This analytically calculated steady-state profile is plotted in Fig. 5*A* (black dots), and is much closer to linear, as opposed to the concave shape obtained in the case of a steady-state PSM length. In fact, it is very slightly *convex* [26]. This almost-linear phase profile also scales with the PSM length in the continuum limit.

**FIG. 5.**
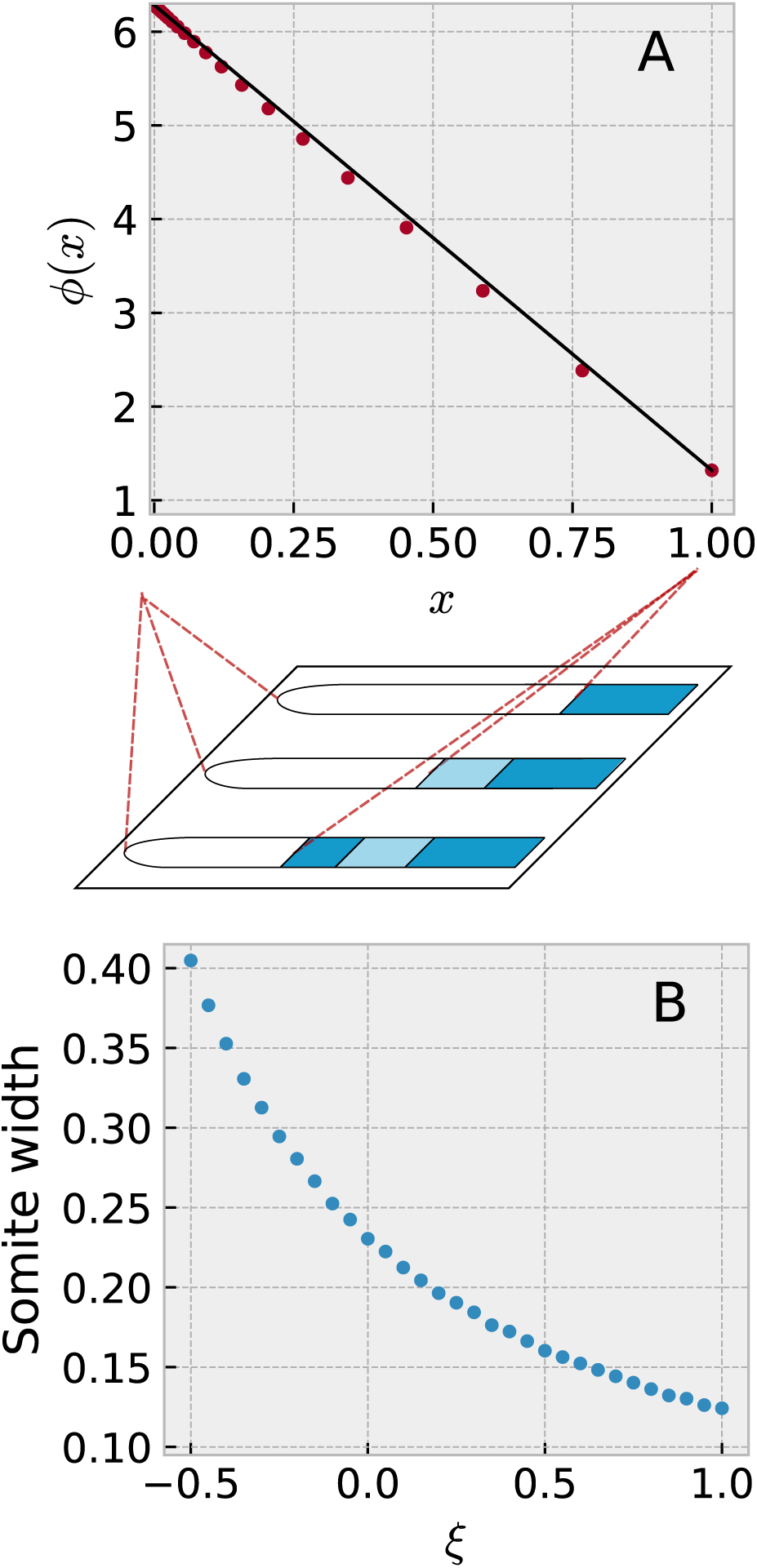
Steady-state phase profile for a PSM that does not grow, but is shortened periodically by removing the anteriormost 0.21 2*π* of phase when the total phase difference between anterior and posterior is 2*π*. The period gradient is linear with *λ* = 0.266. **A** The points show the steady state phase profile just after somite formation, and a (red) straight line between the end points is shown for comparison. The profile is close to linear, and is convex rather than concave in shape. **B** Assuming a perturbation of all periods by an additive amount *ξT*_0_, we plot the actual somite width as a function of perturbation size. The physical size decreases as periods get longer.

In this case too, we can examine the consequence of perturbing all periods in the PSM by a fixed amount *ξT*_0_. Supplementary Section 4 shows the calculation of the new somite widths caused by this perturbation, and Fig. 5*B* plots these as a function of *ξ*. We find that the width decreases as the periods get longer, similar to what we found in the case of constant PSM length. The exponential decrease of PSM length with time and the shift to an almost-linear phase profile are both testable predictions of our model.

## IV. DISCUSSION

The experimental observation in [4] that the total phase difference across ex-vivo mPSMs is 2*π* and that 21% of this phase constitutes the next somite, independent of PSM size, is a curious one. It is not obvious what the consequences of this may be for somites, and even more unclear why it would be necessary or useful (if indeed it is either) for mice embryos to develop in this way. Our work here shows that this observation directly results in a constraint that connects the width of somites and the period gradient across the PSM during somitogenesis. The constraint applies to what we term the phase-width of the somite, while in the particular case where we assume a steady-state PSM length an additional constraint applies to the actual width of the somite. This constraint influences the shape of the phase profile. For a PSM with steady-state length, we predict that the phase profile will be concave, while a PSM with no growth would have an almost linear (slightly convex) phase profile.

The shape of the phase profile is important for at least two reasons. The first is that it affects how travelling waves develop over time – for a concave profile, the waves slow down as they approach the anterior end, while for a convex profile they speed up. The experiments of Lauschke et. al. are in ex-vivo cultures where there is no growth. Our calculations for the no-growth scenario predict an almost linear phase profile, which would predict that the waves propagate with close to constant velocity. This is in fact what [4] observe. In contrast, slowing down of waves, corresponding to a concave profile, is visible in kymographs from zebrafish experiments [3]. The shape of the phase profile thus has a significant effect on the timing of somite formation, and would therefore be worth measuring in more quantitative detail in future experiments.

The second reason is reducing the effect of errors in somite formation. Recent experiments have found that gene-expression noise increases from posterior to anterior [19] in zebrafish. A concave phase profile gets steeper towards the anterior end of the PSM, i.e., the phase difference between neighbouring cells increases from posterior towards the anterior. This diminishes the effect that errors in the phase width have on the actual width of the created somite. The opposite is true for a convex phase profile which flattens out toward the anterior end. These considerations suggest that if somite formation depends on a measurement of phases of the cells, and if, as is likely, these measurements are error-prone, then one should observe smaller errors in the somite widths when the PSM length is steady, compared to later in somitogenesis when it is decreasing.

The constraint of Eq. (3) also has predictable consequences for perturbation experiments, which might be experimentally tractable. One study reduced the number of introns in the Hes7 gene, resulting in more rapid oscillations [27]. They observed shorter segments, i.e., the opposite behaviour of what we expect from our calculations in Section III D based on the experiments in mouse ex-vivo cultures [4]. So it seems that the two experiments contradict each other. The experiment of ref. [27] did not, however, measure the phase difference across the PSM. So, it would be useful to determine whether the assumption of constant phase difference is violated in this case. It would be interesting to study when perturbations of this sort break the assumption of constant difference and when they do not. If perturbations that don’t break the assumption can be found, they would provide a very useful tool to control somite width in a precise and predictable manner.

Another type of perturbation that may be feasible experimentally is to alter the steepness of the period gradient by suitably altering the expression of the morphogen that controls the time period of the somitogenesis clock. In mPSMs if such a perturbation still results in a steady state with a single wave spanning the PSM at any time, then we predict the change in fractional somite width should be close to half the fractional change in the slope of the period gradient (see Fig. 3). Conversely, if the number of waves spanning the PSM increases under this perturbation, then we predict the relationship between the change in the fractional somite width and the change in the slope of the period gradient would become nonlinear.

Our analysis begs the question of how the embryo maintains the constant phase difference across the entire PSM just before each somite formation. Does the embryo “know” that the peak of a travelling wave has reached the anterior end, and send a “signal” to the posterior end to start a new wave? Or is the information transmitted in the other direction, such that the onset of a new peak at the posterior end “causes” the travelling wave to reach the other end at the same time? A third possibility is that this is simply a non-causative correlation caused by some other constraint in the system. We speculate that inter-cellular coupling between the phases of the oscillating cells could be responsible for this behaviour. However, as mentioned before, inter-cellular coupling cannot be too strong or else the cells would start to synchronize despite their intrinsically different time periods, and this has not been observed. It would be interesting to study what kinds of weak coupling in a 1-dimensional line of oscillators with varying time periods could produce travelling waves that are constrained in such a manner. The framework we have introduced here (or the approach of Ares et al. [28], whose model includes coupling which produces synchronized oscillations across the PSM) could be easily extended for this purpose.

These lines of thought also have implications for the mechanisms of somite formation. The well-known clock and wavefront model assumes that somites form when an oscillating cell moves into a sub-threshold region of an existing morphogen gradient that is tied to the growing posterior end of the PSM. Such a model does not necessarily need travelling waves of gene expression, but one could postulate that somites form when the peak of the travelling wave hits some low threshold of the morphogen gradient. Cotterell et al. [8] suggest instead that the somite forms due to reaction-diffusion events in the vicinity of the previous somite when the oncoming travelling wave interacts with a gradient of molecules whose source is the previous somite. It is not clear if there is a simple way to connect such events with the formation of a new wave peak at the posterior end. In both cases, somite formation would be triggered by events at the anterior end and would need some additional mechanism to constrain the total phase difference across the PSM. Recently, a third mechanism has been proposed in mice: Sonnen et al. [9] reported that Wnt and Notch pathways oscillate out-of-phase in cells in the posterior PSM, and in-phase in cells at the segmentation front. They found that the Wnt pathway does not have slow waves travelling periodically from posterior to anterior like the Notch pathway does. Instead, fast-travelling, pulse-like waves were reported[9], which indicates that the Wnt clocks are (nearly) synchronised across the PSM. Thus, with one clock oscillating with frequency dependent on the spatial position of the cell, while the other clock is synchronised (or nearly synchronised) for all cells across the PSM, measuring the phase difference between the two clocks of a single cell would be equivalent to measuring the phase difference between the Notch clock of the posteriormost cell, and the Notch clock of the cell in question, somewhere else in the PSM. This could serve as a signal to trigger somite formation directly dependent on a measurement of the total phase difference across the PSM, a mechanism similar to what was reported by Lauschke et al. [4] and whose consequences we have studied in this paper.

## V. ACKNOWLEDGEMENTS

We are grateful to Alexander Aulehla, Andy Oates and Raj Ladher for discussions. SK thanks the Simons Foundation for funding. JSJ and MHJ acknowledge support from the Danish Council for Independent Research and Danish National Research Foundation through StemPhys Center of Excellence, grant number DNRF116.

## Supplementary Information

### 1. Phase profile in a PSM consisting of a finite number of cells

In this section, we will calculate phase profiles for a system, in which a finite number of oscillators are placed on line. We assume that new oscillators are added at the left end of the line with time intervals *T*_*g*_ = *m*_*L*_*/T*_0_, with *m*_*L*_ *∈* ℕ, *T*_0_ *>* 0, and that the rightmost *m*_*R*_ *∈ ℕ* oscillators are removed with time intervals *T*_*s*_ = *T*_0_ (it is straight forward to substitute another value of *T*_*s*_ in our analysis). Here, *T*_0_ is the time period of the oscillator at the left end of the line, corresponding to the posterior end of the PSM. For simplicity, we furthermore consider the case *m*_*L*_ = *m*_*R*_ = *m*, corresponding to a line in steady state: In a time interval *T*_0_, it grows as much as it is shortened. The system is illustrated in Fig. 1 in the main text. If we choose *t* = 0 to coincide with the addition of an oscillator at the left end of the line, and removal of *m* oscillators at the rightmost part of the line, and assume that the line consists of *N* oscillators at *t* = 0, the number of oscillators on the line is given by

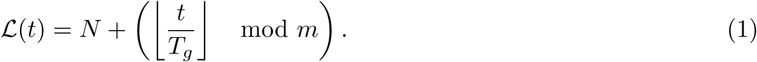

Because the line of oscillators changes its length with time, each oscillator will also change its position on the line, relative to the left end (the time dependent position of an oscillator starting on the leftmost position on the line is shown on the leftmost plot, Fig. 1 A in the main text. Because new oscillators are added on the left end of the line, an oscillator effectively moves one position to the right each time a new oscillator is added. Likewise, every time *m* oscillators are removed from the right end of the line, each oscillator, remaining on the line, effectively moves to the right relative to the length of the line. To formulate an expression for the relative position of an oscillator on the line, let us first consider the case where no oscillators are removed on the right hand side. The line only grows, and it does this by the addition of oscillators on the leftmost end of the line. In this case, no matter the length of the line, the number of oscillators to the right of oscillator *i* is constant. Assuming that the line initially had length *ℒ* (0) = *N,* and that the initial position of the oscillator was 0 *≤ i*_0_ *≤ N −*1, we can exploit this fixed distance to the rightmost end of the line in writing down an expression for the relative position of the oscillator at time *t*,

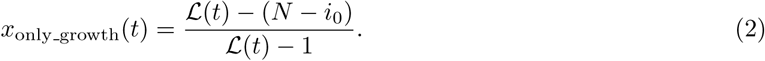

In the above expression, the numerator is an integer, which is *N − i*_0_ smaller than the total number of oscillator on the line. Dividing by the total length (minus 1) yields a number in the interval [0, 1], the relative position of the oscillator. When including periodic removal of oscillator from the right, the expression for the relative position of an oscillator becomes

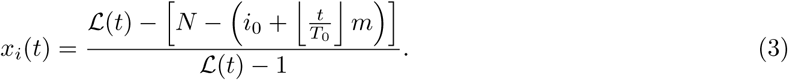

This expression is similar to Eq. (2), except for the term that is added to *i*_0_ in the numerator. This term, 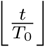 accounts for the removal of *m* oscillators in time intervals of length *T*_0_. This is taken into account because every time oscillators are removed to the right of oscillator *i*, its distance to the rightmost end of the line changes, and this is what we use to calculate *x*_*i*_(*t*).

With the above definitions, we are almost ready to calculate steady state phase profiles. First, however, we must define how oscillation periods change as a function of position on the line of oscillators.

We will assume that oscillation period increases linearly from *T* (*x* = 0) = *T*_0_ at the posterior end (leftmost end of the line, where new oscillators are added) to *T* (*x* = 1) = (1 + *λ*)*T*_0_ at the anterior end (rightmost end of the line, where oscillators are removed from). This assumption is based on experimental observations in mouse mPSMs, as discussed in the main text. *λ* has been determined to be between 0.25 and 0.30 experimentally. Furthermore, we assume that newly added oscillators have initial phase identical to that of the oscillator which occupied the leftmost point on the line until the moment when this new oscillator was added. We implement this assumption by defining the period of oscillators having *negative* spatial positions to have a period and initial phase identical to that of the oscillator on the position *x* = 0 (still referred to as the leftmost oscillator, even though oscillators can now take negative spatial positions, corresponding to oscillators that have not been added to the line yet). Thus, the period distribution is,

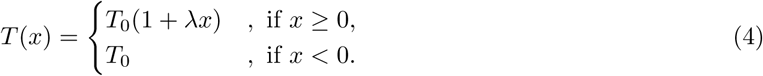

This allows us to write down the phase of oscillator *i* at time *t*,

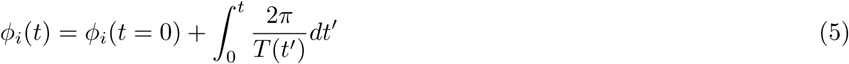

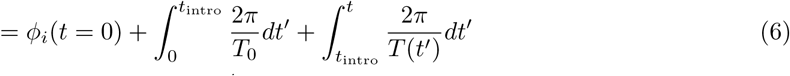

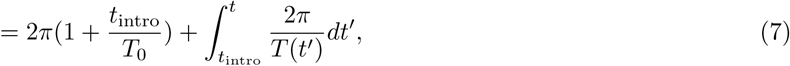

where *t*_intro_ is the time at which the oscillator is introduced at the leftmost end of the line. We can calculate the steady state phase distribution of a line of minimum length *N,* given *T*_*g*_ = *m/T*_0_. We can do this, by identifying *N* oscillators that will take positions 0, 1, …, *N −* 1 at some point, right after oscillators are removed from the rightmost end of the line. First, we notice that all oscillators that end up at the rightmost position, *N −* 1 immediately after removal of oscillators, must previously have taken positions *N −* 1 *− mn, n ∈* ℕ, at corresponding times. One oscillator, which has negative position, but satisfies this, is an oscillator with position *N −* 1 *−* ⌈*N/m*⌉ *m*. If this oscillator starts at position *N −* 1 *−* ⌈*N/m*⌉ *m* at time *t* = 0, it will arrive at position *N −* 1 at time ⌈*N/m*⌉ *T*_0_ immediately after oscillator removal. Another oscillator with initial position *N −* 1 to the left of the initial position stated above will occupy position 0 at the time the oscillator mentioned above occupies position *N*. Hence, by calculating the phase at time ⌈*N/m*⌉ *T*_0_ for all oscillators with initial positions *x*_0_ ∈{*−* ⌈*N/m*⌉ *m −* 1,*−* ⌈*N/m*⌉*m, …, N −* 1 − ⌈*N/m*⌉ *m*]}, we can calculate the steady state phase distribution of the line of *N* oscillators. Without loss of generality, we assume that the leftmost oscillator of the line has phase 2*π* at time *t* = 0, and denote the time at which an oscillator is introduced at the line, *t*_intro_ = *−x*_0_*T*_*g*_ = *|x*_0_*|T*_*g*_. Using Eq. (7),

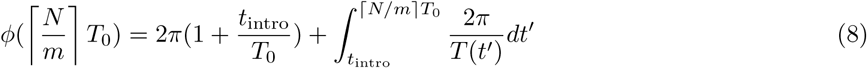

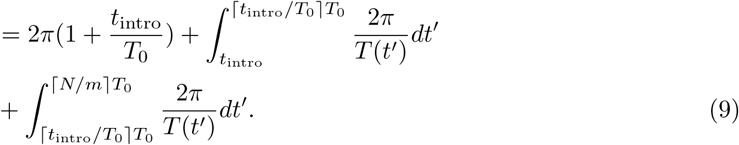

Here we split the last two integrals to make all upper boundaries coincide with oscillator removal. We now convert all integrals to sums over the positions the oscillator takes in each time interval, and insert *t*_intro_ = *|x*_0_*|T*_*g*_. The formula for the phase then becomes

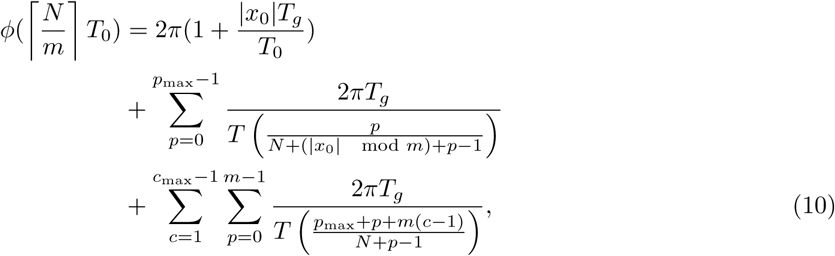

where we have defined *p*_max_ = *m −* (*|x*_0_*|* mod *m*), which is the position an oscillator with initial condition *x*_0_ takes on the line, immediately after oscillators are removed for the first time following its addition to the line, and *c*_max_ = ⌈*N/m*⌉ *−* ⌈*t*_intro_*/T*_0_⌉, which is the number of oscillator removals that an oscillator with initial condition *x*_0_ experiences after being added to the line before *t* = ⌈*N/m*⌉*T*_0_ is reached. In the last term, the first sum takes oscillator removals into account, while the second sum ensures that all *m* positions an oscillator takes between oscillator removals are counted.

### 2. Phase profile in a continuous PSM

In this section, we extend our methods from the previous section to lines of infinitely many oscillators. We will use this method to solve two different example problems. We consider a line of infinitely many oscillators. Oscillators are constantly added on the left end of the line, and a fraction of the line length is removed from the rightmost end of the line in time intervals of *T*_0_. For simplicity, we assume that the length of the line grows linearly between removal of oscillators, and hence, between removals, the length of the oscillator line we define *ℒ* (*t*) = *L*_0_(1 + *βt/T*_0_), where *βL*_0_ is the difference between the maximum and minimum lengths of the PSM. If we now assume that oscillators are removed at times *t* = *nT*_0_ (this amounts to assuming *T*_*s*_ = *T*_0_; once again, it is straight forward to replace this value of *T*_*s*_ with another), *n ∈* ℕ, and that this removal restores the length of the oscillator line to the length it had at *t* = 0, the length is described by

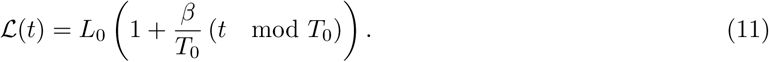

As was the case in our discrete description of the line of oscillators, the position of each oscillator relative to the length of the line, effectively moves right as new oscillators are added at the left end of the line. To express the position of an oscillator as a function of time, we again exploit the fact that the distance between the rightmost point of the line, and an oscillator is constant between removals of oscillators. We write down the relative position of an oscillator as a function of time in the same way as we did in the previous section. First, if no oscillators are removed from the right end of the line, the relative position of an oscillator which had position *x*_0_*L*_0_ (0 *≤ x*_0_ *≤* 1) at *t* = 0, is

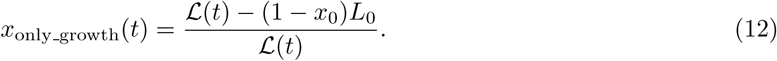

Taking removal of oscillators into account means that *x*_0_ *→ x*_0_ + *β* ⌊*t/T*_0_⌋, since each point moves *βL*_0_ right every time *βL*_0_ is removed from the line from the rightmost end. Therefore,

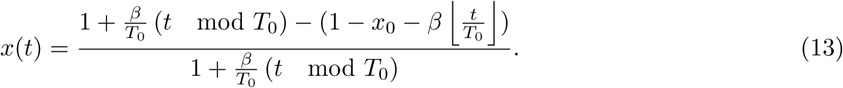

Positions on the line have values *x ∈* [0, 1]. With the same reasoning as in the previous section, for a steady state phase profile, if an oscillator ends up in position *x* = 1 at time *t* = *nT*_0_, *n ∈* ℕ, it has occupied the same positions as all oscillators that ended up at *x* = 1 at *t* = *n*_*−*_*T*_0_, *n*_*−*_ *≤ n −*1. From this follows that these oscillators were added to the line of oscillators at corresponding times between two removals of oscillators. We can characterise such oscillators by the negative position they held at the final oscillator removal before they were added to the line. If the oscillator is added to the line at time 0 *≤ t*_intro_ *≤ T*_0_, this negative position is *x*_start_ = −*βt*_intro_*/T*_0_.

If an oscillator ends up at position *x* = 1 at a time *t* = *nT*_0_, this negative position is

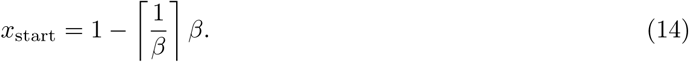

That is, given the period gradient in Eq. (4), and that a newly added oscillator has the same phase as the oscillator to its immediate right, all oscillators that end up at *x* = 1 at a time *t* = *nT*_0_ were added to the leftmost end of the line at time *t*_intro_ = − *x*_start_*T*_0_*/β*, with phase *ϕ*_intro_ = *ϕ*_0_ + *t*_intro_2*π/T*_0_, where *ϕ*_0_ is the phase of the leftmost oscillator of the line right after a removal of oscillators at the right hand end of the line.

We can write down the phase of an oscillator, starting at any position *x*_0_, at any time after it is added to the line,

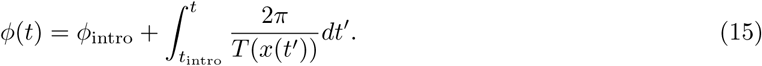

With this formula we can calculate the phase at time *t*, for specific initial conditions *x*(*t* = 0) = *x*_0_. Using this, we can get the steady state phase profile as it looks immediately after oscillator removal, if we calculate *ϕ*(*t*) for a set of initial condition that occupies *x ∈* [0, 1] at a time *t* = *nT*_0_. An interval of initial conditions that satisfies this is *x*_0_ *∈* [*−* ⌈1*/β*⌉ *β,* 1 − ⌈1*/β*⌉ *β*]. The right boundary of this interval, we already concluded will end up at position *x* = 1. It will arrive at position *x* = 1 at time *t* = ⌈1*/β*⌉ *T*_0_, and will eventually be removed from the position at *t* = (⌈1*/β* ⌉+1)*T*_0_. The lower boundary of the interval is exactly *L*_0_ to the left of this point, and hence will be at position *x* = 0, when the rightmost oscillator of the interval arrives at *x* = 1. This reasoning is similar to the one we used in the discrete case. For this reason, we calculate *ϕ*(*t* = ⌈1*/β* ⌉ *T*_0_) for all initial conditions in the mentioned interval, which gives us the phase profile as a function of initial conditions, *x*_0_, right after oscillator removal at time *t* = ⌈1*/β* ⌉ *T*_0_, *ϕ*(*x*_0_, *t* = ⌈1*/β* ⌉ *T*_0_). From this, we obtain the steady state phase profile after oscillator removal *ϕ*(*x*) by replacing *x*_0_ → *x −* ⌈1*/β*⌉ *β*.

For our convenience, we will split the contribution to the phase of the oscillator in question into four parts: 1) The initial phase; 2) Phase acquired before the oscillator is added to the line; 3) Phase that the oscillator acquires after being added to the line, but before the first oscillator removal happens after this; 4) Phase that is acquired at later times. Assuming that all oscillators with negative initial position have initial phase 2*π*, the phase of an oscillator with position *x*(*t* = 0) = *x*_0_ *≤* 0 can then be expressed,

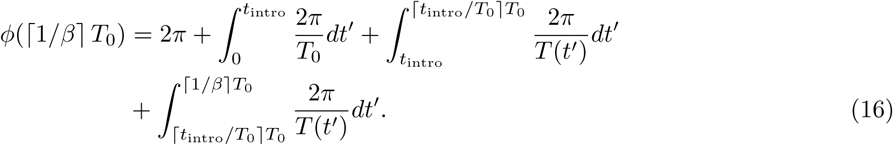

In this expression, the *i*^th^ term corresponds to the *i*^th^ contribution stated above. We will solve the three integrals one by one. The first integral has no explicit time dependence and the solutions is

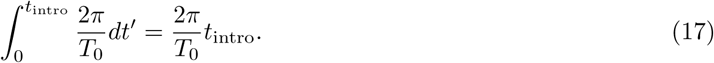

To solve the second integral, we must know *T* (*t* ^*′*^). To know this, we need to know the position of the oscillator as a function of time. We can use the rescaling arguments given above, and write the position for *t*intro *≤ t′ ≤* ⌈ *t*intro*/T*_0_⌉ *T*_0_ as

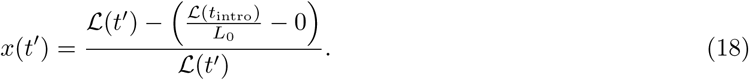

The last oscillator removal prior to *t*_intro_ happens at ⌊*t*_intro_*/T*_0_ ⌋ *T*_0_, and the first oscillator removal following *t*_intro_ happens at ⌈*t*_intro_*/T*_0_⌉ *T*_0_. For this reason, *ℒ* (*t*_intro_) = *L*_0_(1 + (*t*_intro_ − ⌊*t*_intro_*/T*_0_⌋ *T*_0_)*β/T*_0_), and *ℒ* (*t′*) = *L*_0_(1 + (*t −* ⌊*t*_intro_*/T*_0_⌋ *T*_0_)*β/T*_0_). Inserting this in the expression above, we get

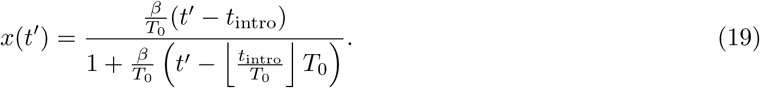

We can insert this in the expression for the period as a function of position in Eq. (4), and insert in the second integral above. We get

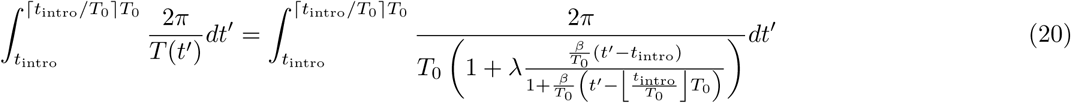

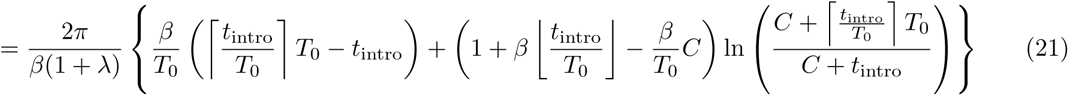

with

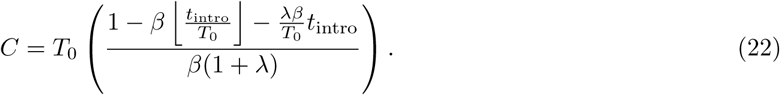

Having solved this integral, we now turn to the last integral in the expression of the phase at time ⌈1*/β* ⌉*T*_0_. The final integral represents the phase that the oscillator acquires after the first oscillator removal. This time interval is an integer number of periods of the leftmost oscillator. The number of periods that the leftmost oscillator goes through before the time *t* = ⌈1*/β* ⌉ *T*_0_ is reached, we denote *c*_max_ = ⌈1*/β*⌉ *−* ⌈*t*_intro_*/T*_0_⌉. From this insight, we can write the integral as a sum *c*_max_ integrals over time intervals of length *T*_0_. This is expressed as follows,

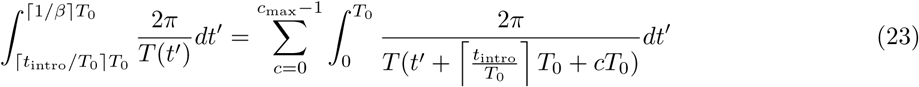

To solve this we once again need to know the length of the line at the time at which the integrand is evaluated. In 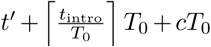 the final two terms are both integer powers of *T*_0_. Since *ℒ* (*t*) = *ℒ* (*t* + *jT*_0_), *j ∈* ℕ, we know that 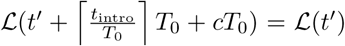. To write down the position of the oscillator at time 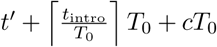, we need to know the position of the oscillator at time 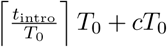. After each full time interval *T*_0_, the line is restored to length *L*_0_, and all oscillators have moved *βL*_0_ to the right. For this reason, at time 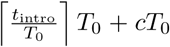, an oscillator with initial condition *x*_0_ has position 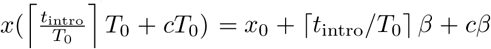 We can use these pieces of information to express the position of the oscillator at time 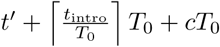 as follows

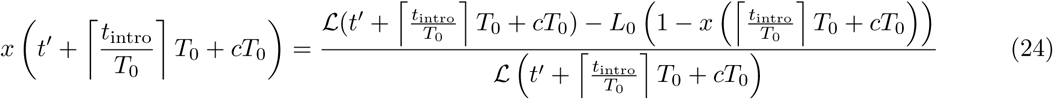

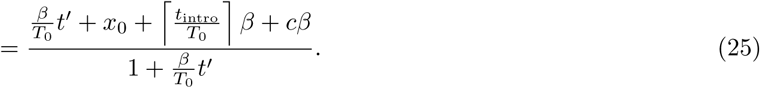

This, we can insert into Eq. (4) to get the oscillator period at the time in question. The final integral then becomes

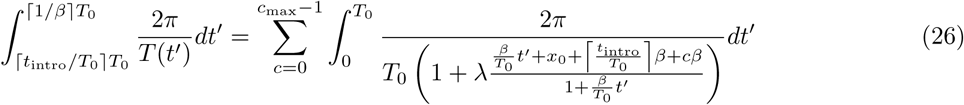

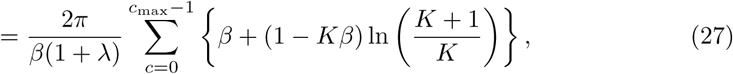

where

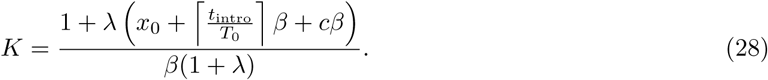

The steady-state phase profile is obtained by adding all 4 contributions, and this is the black curved plotted in Fig. 2 in the main text.

### 3. Phase profile in a PSM that does not grow

In the previous sections we analyzed a line of oscillators that grows as much as it is shortened in one posterior period. We now turn to another important special case: A line in which there is no growth, only oscillator removal. In this case, the length of the line is conserved between oscillator removals, and an oscillator does not change its position relative to line length between oscillator removals. We will assume that 1) the phase difference between the two ends of the line is Φ_before_ at oscillator removal (Inserting Φ= 2*π* is the special case of mouse mPSMs with no growth); 2) That a phase profile is rescaled to line length but otherwise identical (mod 2*π*) at oscillator removal; 3) That oscillation period increases linearly along the line like above; 4) That *T*_*s*_ = *T*_0_ (once again, it is straight forward to substitute this assumption with another value of *T*_*s*_; 5) That the 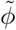 rightmost phase is removed at oscillator removal – this means that a certain fraction of the line length 1 − *x*_*c*_ is removed from the right end of the line at oscillator removal (that is, the oscillator with position *x*_*c*_ *∈* (0, 1) before oscillator removal has position *x* = 1 after oscillator removal). In assumption 5), the value of 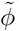 is intimately connected to the period-profile on the line; we will be using the experimentally observed value 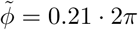. With these assumption we will estimate the phase profile over the line and determine *x*_*c*_, or equivalently the fraction of the oscillator population that is removed at each oscillator removal 1 - *x*_*c*_.

Assumption 5) above means that an oscillator which has position *xx*_*c*_ just before oscillator removal will have position *x* until next time oscillators are removed. The oscillator changes its phase *δϕ*(*x*) = 2*πT*_0_*/T* (*x*) between these two consecutive oscillator removals. But because the leftmost oscillator acquires 2*π* of phase between consecutive oscillator removals, and the phase profile is in steady state, *ϕ*(*x, t* = *nT*_0_) = 2*π* +*ϕ*(*x, t* = (*n* + 1)*T*_0_), *n ∈* ℕ. These insights make us capable of writing down the following equation for the phase of the oscillator on position *x* just before oscillator removal

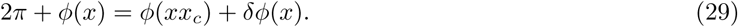

In accordance with the assumptions above, we, without loss of generality, take the phase in the line endpoints to be *ϕ*(*x* = 0) = Φ_before_ and *ϕ*(*x* = 1) = 0 just before oscillator removal. We now work towards an expressions that will allow us to determine the phase in infinitely many different points, and determine *x*_*c*_ under our above assumptions. Evaluating Eq. (29) in 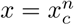 yields

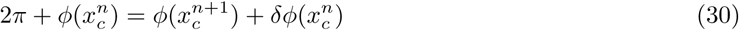

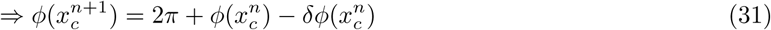

This is a recursive relation. Given a period distribution, which in this case is *T* (*x*) = *T*_0_(1 + *λx*), we can use this to determine *x*_*c*_ such that Φ_before_ and 0 are the phases of the endpoints of the line (these are specific to our example biological system and could be chosen differently if wanted). We can now insert *δϕ*(*x*) = 2*πT*_0_/(1 + *λx*), and reduce the expression in Eq. (31) to obtain the recursive relation

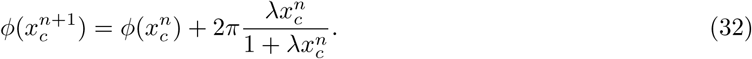

Using 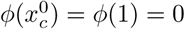, and the recursive relation above, the phase at any point 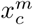 can be calculated

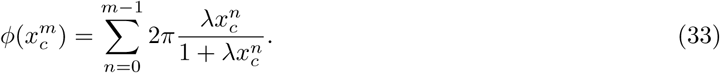

Taking the limit *m → ∞*, we know 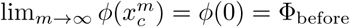, and this gives us

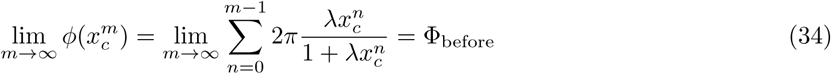

This is an equation in *x*_*c*_, which is nontrivial to solve analytically. Numerically, we evaluate the sum to e.g. *n* = 550 for different values of *x*_*c*_. For Φ = 2*π*, we find that *x*_*c*_ = 0.767622 solves the equation. A bound for the error on this evaluation can be found by the following estimation

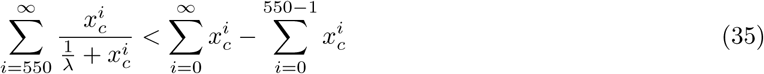

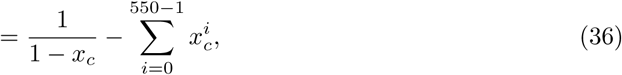

where we used that 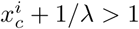, and 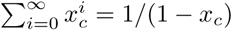. This yields an error on the evaluation smaller than 10^-15^.

Having estimated *x*_*c*_ = 0.767622, we now know the fraction of oscillators that are removed periodically, 1*−x*_*c*_ = 0.232378, and can plug *x*_*c*_ = 0.767622 into Eq. (33) to obtain the steady state phase in any point 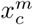, *m ∈* ℕ. In the previous sections, we plotted phase profiles *after* oscillator removal, and not *before* oscillator removal like here. The steady state phase profile after oscillator removal is obtained by replacing *x*→ *x/x*_*c*_ for the evaluated points, and removing the point *x* = 1. This is plotted in Fig. 5A in the main text.

### 4. Changing somite width in a PSM that does not grow

In the previous section, we determined the somite width in a PSM that does not grow. This we did for a specific value of *λ*. In this section, we imagine perturbing the period gradient such that every oscillator has its period altered by an additive amount *ξT*_0_. So the new period distribution is *T* (*x, ξ*) = *T*_0_(1 + *xλ* + *ξ*). We assume that a somite is formed once every posterior period, and we assume that the phase width of the somite is equal to the phase difference that occurs between posterior and anterior in one posterior period. Lastly, we assume that the phase difference between anterior and posterior is 2*π* at the time of somite formation, and that somites form with period *T*_*s*_ = *T*_0_. It is straight forward to substitute the assumed values of Φ_before_ and *T*_*s*_ with other values.

Eq. (29) is still valid, except that we now have an additional variable,

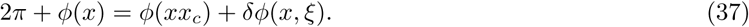

*δϕ*(*x, ξ*) is equal to the difference that occurs between posterior and anterior in one posterior period, and is given by

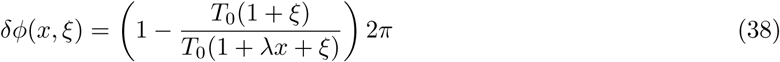

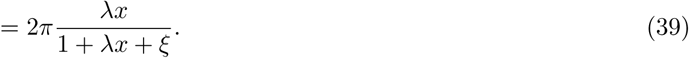

We now proceed as we did in the previous section. Inserting 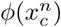 recursively gives us the equation

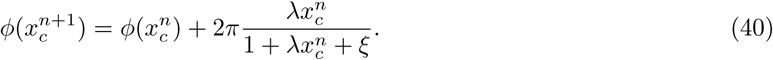

If we use 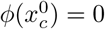, we can calculate

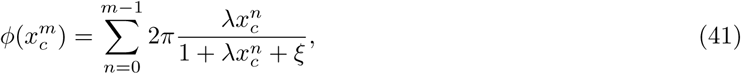

and with this, we can demand that the posteriormost oscillator has phase Φ_before_ at somite formation,

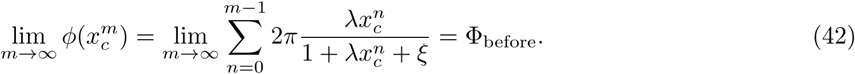

So for a given *ξ*, the corresponding *x*_*c*_ satisfies the equation

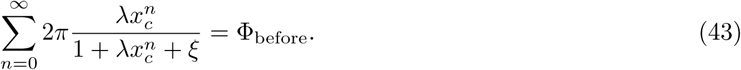

We evaluate the first 550 terms of this sum, for Φ_before_ = 2*π*, and use this to find the *x*_*c*_ that satisfies Eq. (43). The error on this evaluation is estimated as we did in the previous section,

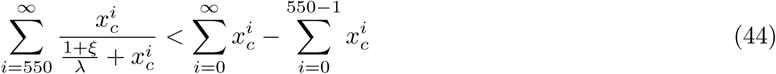

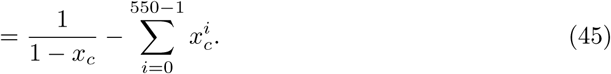

This holds if (1 + *ξ*)*/λ >* 1. This is definitely true for *ξ ∈* [−0.5, 1], which is the range that we plot *x*_*c*_(*ξ*) for in Fig.5B in the main text. In the main text we refer to *x*_*c*_(*ξ*) as the physical somite width.

### 5. Physical somite width as a function of period perturbation size

The calculation in Supplementary Section 2 gave us the phase profile in a steady-state PSM whose maximal length is *βL*_0_ longer than its minimal length *L*_0_. In this calculation, we have not assumed a specific Φ_before_. For different choices of *β* and *λ*, we get different values for the phase difference Φ_before_, and phase width 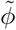. This lets us examine how perturbing the period for all cells in the PSM by an amount *ξT*_0_ changes the physical somite width. We will do this with the following approach

1. Choose a parameterx *β*,
2. Try different input values of *λ*, and find the value that corresponds to the wanted Φ_before_, e.g. Φ_before_ = 2*π*,
3. Calculate what value of *ξ*, corresponds to the determined value of *λ*.

The first two steps in this approach are straight forward to carry out, using the analytical expression for the phase profile in a steady-state PSM in the continuum limit. We only need to figure out how to convert a chosen *λ*-parameter to a *ξ*-value. First, we note that if oscillators are removed once every posterior period (assuming *T*_*s*_ = *T*_0_, other values for *T*_*s*_ are easy to plug in to the calculations), the phase width of a somite in a PSM with posterior period *T*_min_ and anterior period *T*_max_ is given by,

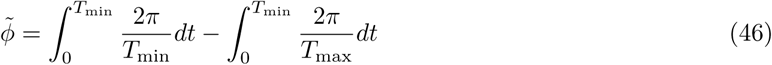

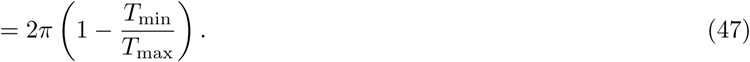

So the phase width is not determined by the *difference* between the posterior and anterior periods, but by the *ratio* between the posterior and anterior periods. We will now take advantage of our theoretical framework from Section 2 of this supplementary file. In our framework, we can choose any value for the difference in periods over the PSM, *λ*, we like. However, suppose that we know that only a single value *λ* = *λ*_0_:= 0.21/0.79 corresponds to the period gradient in the PSMs observed in an *unperturbed* experiment. Suppose all other values of *λ* correspond to the period gradient in experiments where an additive perturbation *ξT*_0_ affects all cells in the PSM. In the rest of this section, this is what we will assume. By assuming this, we will provide a formula for matching the chosen *λ* to a unique value of the perturbation size, *ξ*, given *λ*_0_.

For a chosen parameter *λ*, the ratio between periods in the simulation is

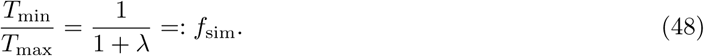

As described above, we now assume that any *T*_min_ from our simulations can be written *T*_min_ = *T*_0_(1 + *ξ*), and likewise for *T*_max_ = *T*_0_(1 + *λ*_0_ + *ξ*). If we demand that the observed ratio between periods *f*_sim_ is equal to the ratio between these perturbed periods, we get

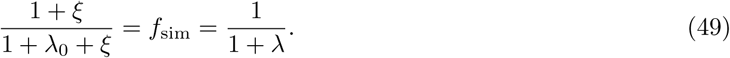

Solving for *ξ* gives us

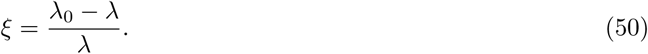

This formula lets us match the *λ*-value, which ensures the wanted Φ_before_ for a chosen *β*, with a *ξ*-value. We plot matching *β* and *ξ*-values for Φ_before_ = 2*π* in Fig. 4B in the main text.

### 6. Phase profile shape and somite size for different growth conditions

In the previous sections, we have examined somite size, and phase profiles in the absence of growth in PSMs, and in PSMs with steady-state length. We found that while the phase profile was convex in the absence of growth, it was concave in PSMs with steady-state lengths. We also found that the physical somite size was larger in PSMs that do not grow. In this section, we will use perturbation theory to argue

- For any *T* (*x*), *x ∈* [0, 1], which is an increasing function of *x*, and PSM that has steady-state length, the corresponding steady-state phase distribution is concave.
- How the steady-state phase distribution could become convex, if the PSM does not have steady-state length.

#### 6.1 Concave phase profiles if PSM length is in steady state

First, we examine the phase profile in a PSM in steady state. Suppose that the PSM consists of a larger number of cells. We use the continuous variable *x ∈* [0, 1] to describe the position of each cell relative to the posterior end (at *x* = 0) and the anterior end (at *x* = 1). Let *T* (*x*) be the period gradient of the PSM, and let this be increasing from posterior to anterior. Suppose that two cells have initial positions *x*(*t* = 0)_first_:= *x*_0,first_ equal to *x*_0_ = *x*^*\**^, and *x*_0,second_ = *x*^*\**^ +*ϵ*, where 0 ≤ *x*^*\**^ *<* 1, and 0 *< ϵ ≪*1. Let us assume *t* = 0 to be immediately after somite formation, and let the phase difference between the two cells be *δϕ*_*ϵ*_ = *ϕ*(*x*) − *ϕ*(*x* +ϵ) *>* 0. We now examine how the phase difference between these cells changes between *t* = 0, and just after the following somite formation at *t* = *T*_*s*_. The change in phase difference between the two cells in this time period is

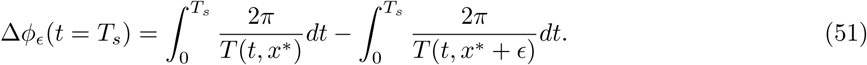

Now, since *ϵ ≪* 1, we expand the fraction

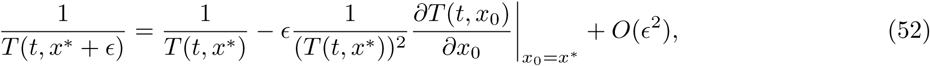

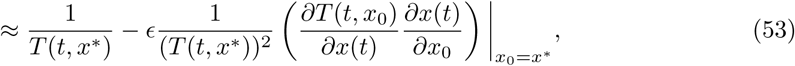

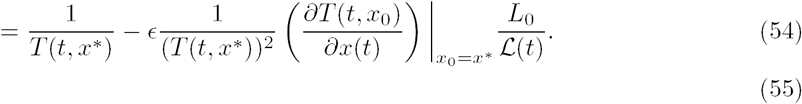

Here we used Eq. (12) to calculate *∂x*(*t*)*/∂x*_0_. Inserting this expression in Eq. (51) yields,

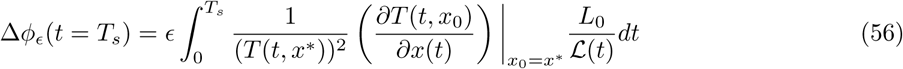

Since *T* (*x*) is increasing and positive, and since *ℒ* (*t*) is positive and increasing between somite formation, Δ*ϕ*_*ϵ*_ *>* 0. This means that the phase difference between the two cells increases between the two somite formations. The phase difference is the same after the somite formation at *t* = *T*_0_, and because the PSM length is in steady state, the difference in position between the two cells is still *ϵ* at *t* = *T*_0_. We can determine whether the phase profile is convex or concave by comparing whether the phase profile is decreasing more quickly at positions that are more posterior or more anterior. This tells us whether the phase profile is convex or concave because a decreasing, concave function has a negative second derivative, while the second derivative is positive for a decreasing, convex function (See Fig. 1S in this Supplementary file). The phase profile gradient between the cells at their initial position is *δϕ*_*ϵ*_*/ϵ*, and the phase profile gradient between the cells at their final position is (*δϕ* + Δ*ϕ*_*ϵ*_)*/ϵ*. Calculating the ratio yields

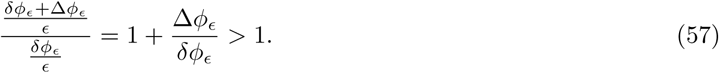

From this we conclude that the steady-state phase profile decreases faster as *x* is increased. Or equivalently: *the steady-state phase profile is concave*.

#### 6.2 Convex phase profile with *T* (*x*) linear, increasing

In the previous subsection we found that the phase difference between two cells increases between somite formation if *T* (*x*) is an increasing function. This was expressed in Eq. (51). With this in mind, one may wonder how one can obtain a convex phase profile with an increasing period profile *T* (*x*), as we found in the case of no growth. The answer lies in the shortening of the PSM: Even though the phase difference between two cells increases between somite formations, the somite formation itself causes the difference in position of the cells to grow relative to the length of the PSM. This may influence the ratio between the phase profile gradients greatly.

**Figure 1S:**
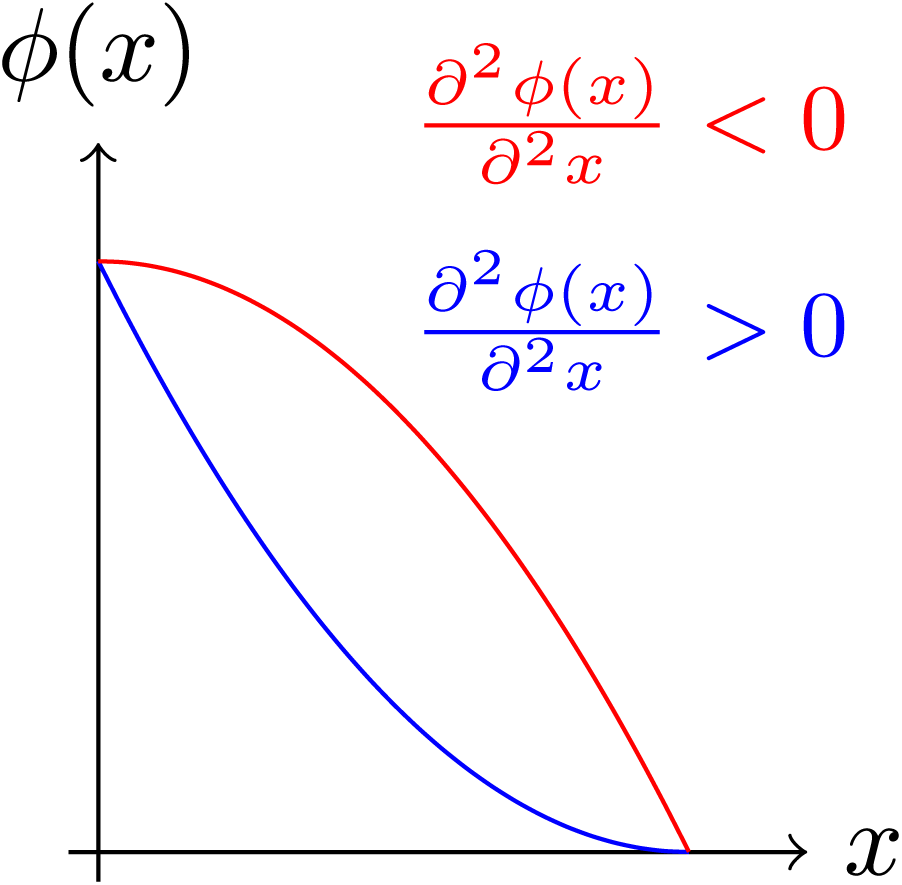
Illustration of decreasing functions that are concave (red), and convex (blue). The concave function has a negative second derivative, while the convex function has a positive second derivative.

Let us return to the case of a PSM with period-profile *T* (*x*) = *T*_0_(1 + *xλ*), that does not grow. We can evaluate Eq. (51) in this case,

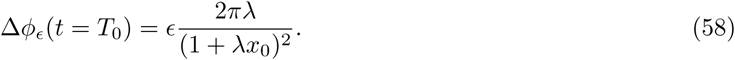

If two cells were a distance *ϵ* apart from each other before somite formation, and if *x*_*c*_ is the length of the PSM after somite formation, the distance between the same cells after the somite formation will be *ϵ/x*_*c*_ relative to the new PSM length. Taking this into account, we can now calculate the ratio between the phase profile gradient between two cells at two consecutive somite formation events, as we did in the previous subsection,

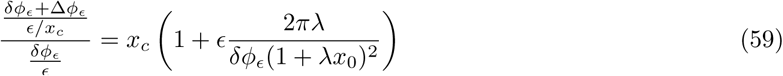

if *x*_*c*_, and *ϵ/δϕ*_*ϵ*_ are sufficiently small, this may be less than 1, resulting in a convex phase profile. This is the case for the phase profile plotted in Fig. 5A in the main text.

